# Microtubule organization within mitotic spindles revealed by serial block face scanning EM and image analysis

**DOI:** 10.1101/087866

**Authors:** Faye M. Nixon, Thomas R. Honnor, Georgina P. Starling, Alison J. Beckett, Adam M. Johansen, Julia A. Brettschneider, Ian A. Prior, Stephen J. Royle

**Author notes:** joint first author (Faye M. Nixon, Thomas R. Honnor).

## Abstract

Serial block face scanning electron microscopy (SBF-SEM) is a powerful method to analyze cells in 3D. Here, working at the resolution limit of the method, we describe a correlative light-SBF-SEM workflow to resolve microtubules of the mitotic spindle in human cells. We present three examples of uses for this workflow which are not practical by light microscopy and/or TEM. First, distinguishing closely associated microtubules within K-fibers; second, resolving bridging fibers in the mitotic spindle; third, visualizing membranes in mitotic cells, relative to the spindle apparatus. Our workflow also includes new computational tools for exploring the spatial arrangement of MTs within the mitotic spindle. We use these tools to show that microtubule order in mitotic spindles is sensitive to the level of TACC3 on the spindle.

## 1 Introduction

In readiness for cell division, eukaryotic cells build a mitotic spindle. This miniature machine, made of microtubules (MTs) and associated proteins, maneuvers the duplicated chromosomes and segregates them so that the two daughter cells each receive a complete copy of the genome. Understanding how this MT-based structure is organized and how it works is a major theme in cell biology (Petry, 2016).

In human cells, the mitotic spindle is a fusiform structure, which at metaphase is virtually spherical. MTs radiate from the two spindle poles, in one of three classes. 1) Astral MTs extend from the poles back toward the plasma membrane; 2) Interpolar MTs run from one pole toward the other pole;3) Kinetochore MTs which are attached to the kinetochore on the chromosome (Mastronarde et al., 1993; McDonald et al., 1992; Helmke et al., 2013). Kinetochore MTs are bundled together to form kinetochore fibers (K-fibers) and at metaphase, 20–40 MTs constitute a single K-fiber bundle (Booth et al., 2011, 2013; Sikirzhytski et al., 2014; McEwen et al., 1997). A network of crosslinking material, termed the “mesh”, is thought to hold kinetochore MTs together throughout their length (Hepler et al., 1970; Nixon et al., 2015). So far one protein complex containing TACC3, clathrin and ch-TOG has been identified as being a component of the mesh (Booth et al., 2011; Nixon et al., 2015).

The MTs of the mitotic spindle are densely packed and so resolving their organization in 3D is challenging. Advances in super-resolution imaging and expansion microscopy have improved the view of mitotic MTs by light microscopy (Mikhaylova et al., 2015; Chozinski et al., 2016), yet the gold standard method is still electron microscopy (EM). Reconstructions of entire spindles in 3D using serial sectioning and transmission electron microscopy (TEM) have told us much about mitotic spindle structure (Mastronarde et al., 1993; McDonald et al., 1992; Redemann et al., 2016). However these methods are extremely laborious, technically challenging, and prone to error. The advent of automated EM methods, such as serial block face scanning electron microscopy (SBF-SEM) has promised to speed up the analysis of complex cellular structures in 3D (Hughes et al., 2014). The spatial resolution of SBF-SEM is reported to be around 10nm, making it ideal for the study of MTs (25nm diam.) in the mitotic spindle. Whether this resolution can be reached routinely for such analysis is unclear.

The first aim of the present study was to apply SBF-SEM to the problem of resolving MTs in the mitotic spindle. The resulting datasets are large and require robust statistical methods for analysis and our second aim was therefore to provide computational tools to aid understanding of such datasets. The resulting workflow was then used to further understand how TACC3 levels influence MT organization in mitotic spindles.

## 2 Results

### 2.1 Visualization and 3D reconstruction of mitotic spindle MTs

We began by finding the optimal conditions for imaging MTs in mitotic cells by SBF-SEM. These conditions, described in the Methods section, allowed us to image a volume containing almost the entire spindle and to visualize bundles of MTs in the spindle running from the pole to the chromosomes (Figure 1A). The resulting datasets could be segmented, rendered in 3D, and analyzed computationally. This workflow resulted in the visualization of a 3D rendering of the mitotic spindle (Figure 1B). The architecture of the mitotic spindle was visible at sufficient resolution to discern individual K-fibers, interpolar MTs (ipMTs), and in some instances, kinetochores (Figure 1C and Supplementary Movie 1). Under these optimized conditions, there were significant limitations. The spatial resolution we achieved was probably not sufficient to discern individual MTs. For example, almost no astral MTs could be seen in any of the SBF-SEM datasets. Therefore all the data presented below likely come from segmentation of at least two MTs in a single location. Moreover, the voxel size of 12 x 12 x 60nm meant that the resolution in the z-dimension was less than that in x and y. Despite this, our simple and rapid workflow allows the accurate mapping of mitotic spindle architecture sampled beyond the resolution limit of light microscopy.

### 2.2 Spatial statistics of MTs visualized by SBF-SEM

The large-scale high-resolution view of MTs resulting from our workflow was well-suited for the development of geometric models to describe mitotic spindle organization. These methods and associated code are detailed in the Methods section, and an overview is presented here.

**Figure 1:**
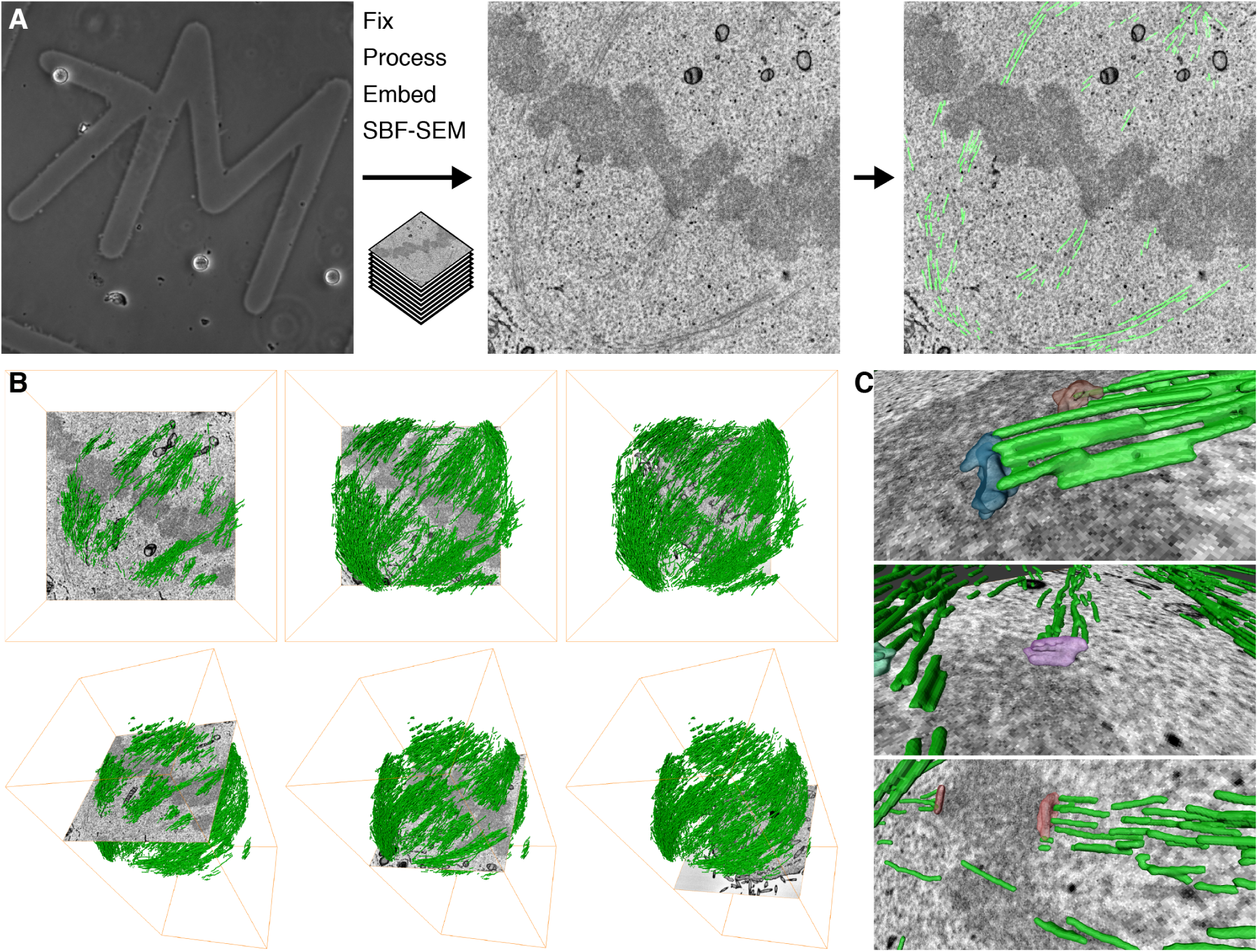
SBF-SEM of mitotic spindles. (**A**) Practical workflow. A single mitotic cell on a gridded dish is first visualized by light microscopy (left) to assess its mitotic stage and transfection status. The cell is then fixed and embedded for SBF-SEM. A series of SEM images are captured at each section through the block (typical image is shown, center). Using Amira, the MTs are segmented along with any other cellular features, as required (right). (**B**) 3D rendering of MTs in the mitotic spindle is shown with an orthoslice at three different depths (left to right), *en face* (above) or as a tilted view (below). **(C**) 3D rendering of example kinetochores and their associated K-fibers (green).

First, the volumetric density of MTs in the spindle could be determined. As an input, we take the segmented MTs from SBF-SEM data and this is used to calculate the total volume of MTs. This value is then normalized to the volume of spindle which is visible in the dataset (Figure 2A). This normalization step is essential because at the magnification necessary to resolve MTs, the spindle volume is often incomplete. To test the sensitivity of this volume density calculation, we compared cells treated with cold (4°C) *versus* warm (37°C) media. When cells are treated with cold media, any MTs which are not stably attached at both ends become depolymerized leaving only the K-fibers intact (Rieder, 1981). In HeLa cells and in two different cell lines derived from Glioblastoma patients, we measured a decrease in MT density to approximately one-third of the volume recorded in control cells, in warm media (Figure 2B).

Second, the spatial organization of MTs in the spindle is analyzed. The aim is to determine the order – or degree of alignment – of MTs in the mitotic spindle. To do this, an idealized spindle is generated computationally using the coordinates of the segmented MTs, and simple geometric principles (Figure 3 and Methods section 4.4). The measured angle between each MT segment and its idealized counterpart is calculated (*θ*^(*i*)^ in Figure 3A). For an ideal mitotic spindle, in which all microtubules are aligned, all angles will equal 0°. For real mitotic spindles, the angles deviate considerably from the ideal. Therefore, the magnitude of the angle deviations from the idealized spindle indicate a departure from perfect alignment. For example, the microtubules in the control GFP-expressing cell shown throughout this paper, have a median angle of 11.2° (IQR = 14.8, 5.2 – 20.0°) (Figure 4A). Previously, we found that overexpression of TACC3 caused changes in the degree of alignment of MTs (Nixon et al., 2015). These conclusions were based on tomography of a single thin section cut orthogonally to the K-fiber axis. Our new statistical method meant that this comparison could now be done on a near-global, whole-spindle scale. To look deeper into this phenomenon, we wanted to increase or decrease the TACC3 levels on the spindle and examine changes in MT order. Therefore, in addition to GFP-expressing control cells, we processed cells conditionally overexpressing GFP-TACC3 and also cells where the endogenous TACC3 was depleted by RNAi and a mutated form (S558A) of GFP-TACC3 that was refractory to RNAi was expressed. This mutant cannot bind clathrin and therefore cannot localize to the mitotic spindle (Hood et al., 2013). These three genotypes were examined in both warm and cold conditions (Figure 4C). Deviations from the ideal were compared using Π (Θ_1_, Θ_2_) for all combinations of data (see Methods section 4.4.5 for further details). This comparison of angle deviations from these datasets showed that mitotic spindles in cells with more TACC3 had less order than controls, under warm conditions (Figure 4A and 4C). Similar deviations were seen in the cells expressing TACC3(S558A) in place of endogenous TACC3. This analysis extends, and is in agreement with, our previous conclusion that MT order in mitotic spindles is influenced by TACC3 levels on the mitotic spindle (Nixon et al., 2015). The comparison between warm- and cold-treated samples revealed that MTs in cells expressing TACC3(S558A) in place of endogenous TACC3 were more disorganized after cold treatment. Interestingly, the cold-treated cells overexpressing TACC3 did not show the same phenotype. The order of MTs in this condition was similar to cold-treated GFP-expressing cells, which may suggest that elevated TACC3 levels protects MTs against cold-induced changes in spatial organization.

**Figure 2:**
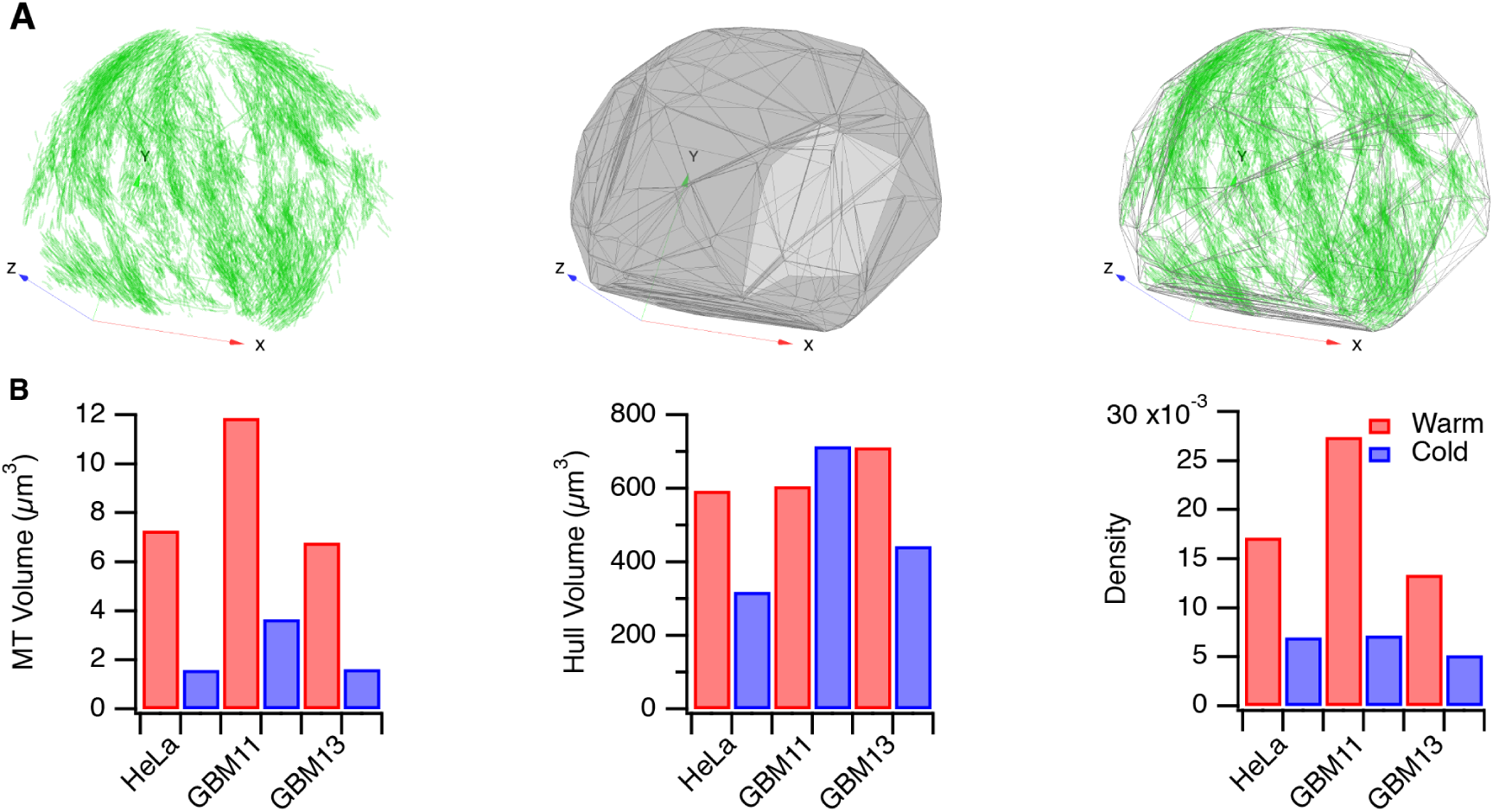
Volume calculations for mitotic spindles. (**A**) Map of MTs in a HeLa cell at metaphase. A 3D convex hull is drawn around all points in the map and used for volume calculation and normalization of MTs into a density. The same cell is shown in Figure 1. (**B**) Plots of MT volume, Hull volume and Density (MT volume/Hull volume) for three different cell types. A comparison of warm (red) versus cold (blue) is shown. *N*_cell_ = 1 per condition.

**Figure 3:**
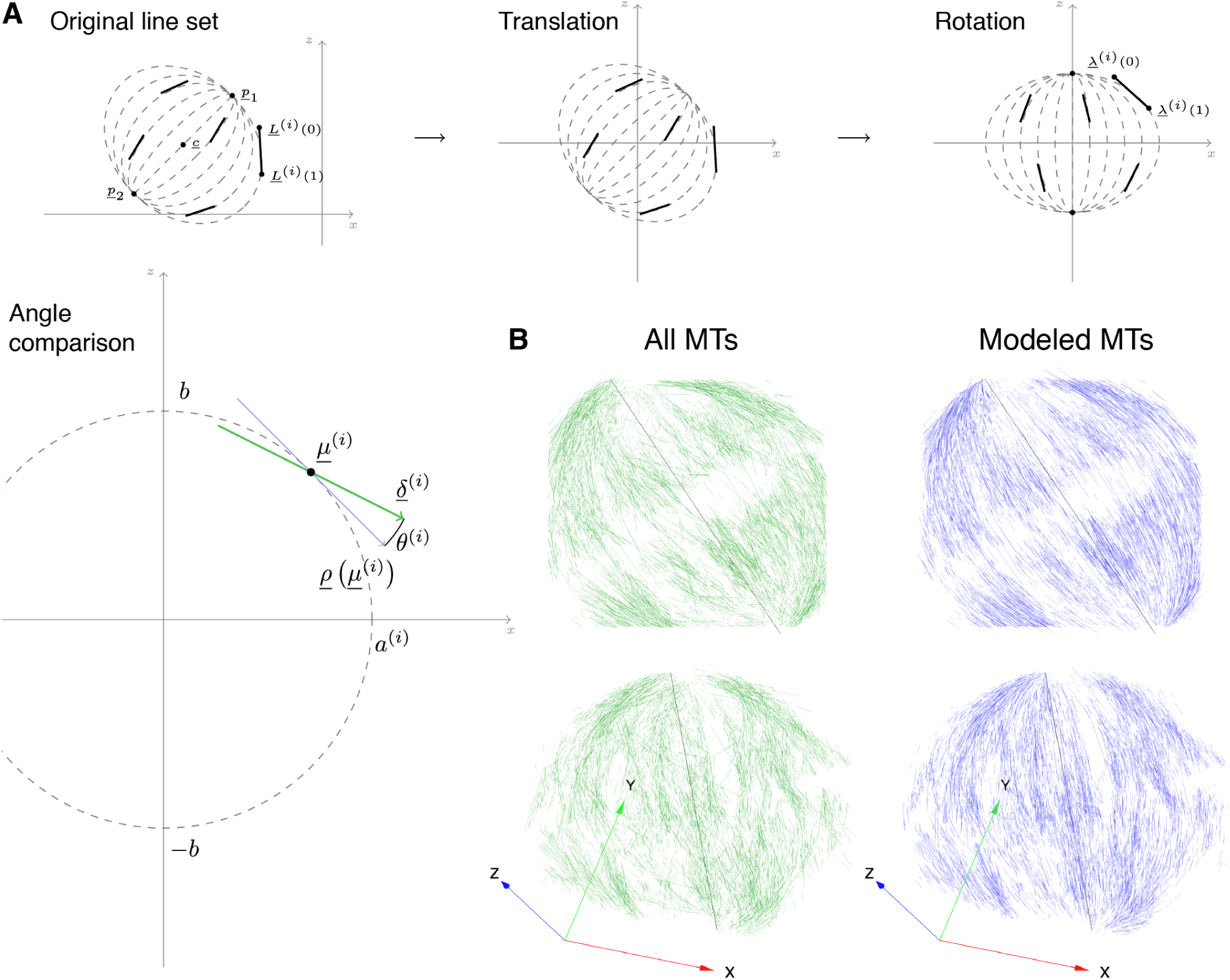
Using a simple geometric model to obtain idealized mitotic spindles from measurement data. (**A**) Schematic diagrams to illustrate how the idealized mitotic spindles were generated. The real data set is translated and rotated so that the spindle is stood up, with the spindle poles aligned on the z-axis at *x* = *y* = 0. Ellipsoids which pass through the spindle poles are generated for the midpoint of each MT and the tangent is used to generate the idealized MT counterpart. The angle between the real and ideal line is recorded for each MT. (**B**) An example of an idealized model spindle (blue, right) was created from the MT positions in a real data set (green, left). The spindle is shown *en face* (XY view, above) and tilted (below). Deviations of the real data from the model spindle were measured by taking the angle of real MTs *versus* their idealized counterpart. The cell from which this data derives is shown in Figure 1.

### 2.3 Two further observations from SBF-SEM: spindle encapsulation and bridging fibers

In addition to the mitotic spindle and associated chromosomes, the membranes of mitotic cells could also be rendered in 3D. This allowed us to explore the relationship between intracellular membranes and the mitotic spindle. We found that the mitotic spindle was situated within an “exclusion zone” which was largely free of membranes and organelles (Figure 5, Supplementary Figure 1 and Supplementary Movie 2). Outside of this zone, the endoplasmic reticulum (ER) was densely packed and mitochondria were distributed within the folds of ER. These observations agree with previous work which described the organization of ER membranes in mitotic cells (Puhka et al., 2007, 2012) and with more recent work which proposed that although mitosis in human cells is open, the spindle is encapsulated by membrane (Schweizer et al., 2015). The purpose of this encapsulation is to exclude organelles and concentrate the proteins necessary to build a mitotic spindle rapidly (Schweizer et al., 2015).

Whilst studying the organization of MTs in the mitotic spindle, we also had the opportunity to look for the presence of “bridging fibers”. These small bundles of MTs have been proposed to span from one K-fiber emanating from one pole, across the sister kinetochores and reach the K-fiber which emanates from the opposing pole (Kajtez et al., 2016; Tolic and Pavin, 2016). Close inspection of our segmented datasets showed the presence of structures which may correspond to bridging fibers. Examples are shown in Supplementary Movie 3 and a single example is shown in Figure 6. We could not find evidence for bridging fibers associated with every K-fiber pair, although this is probably because our coverage is incomplete rather than an indication that there is not a one-to-one association of bridging fibers with each K-fiber pair.

**Figure 4:**
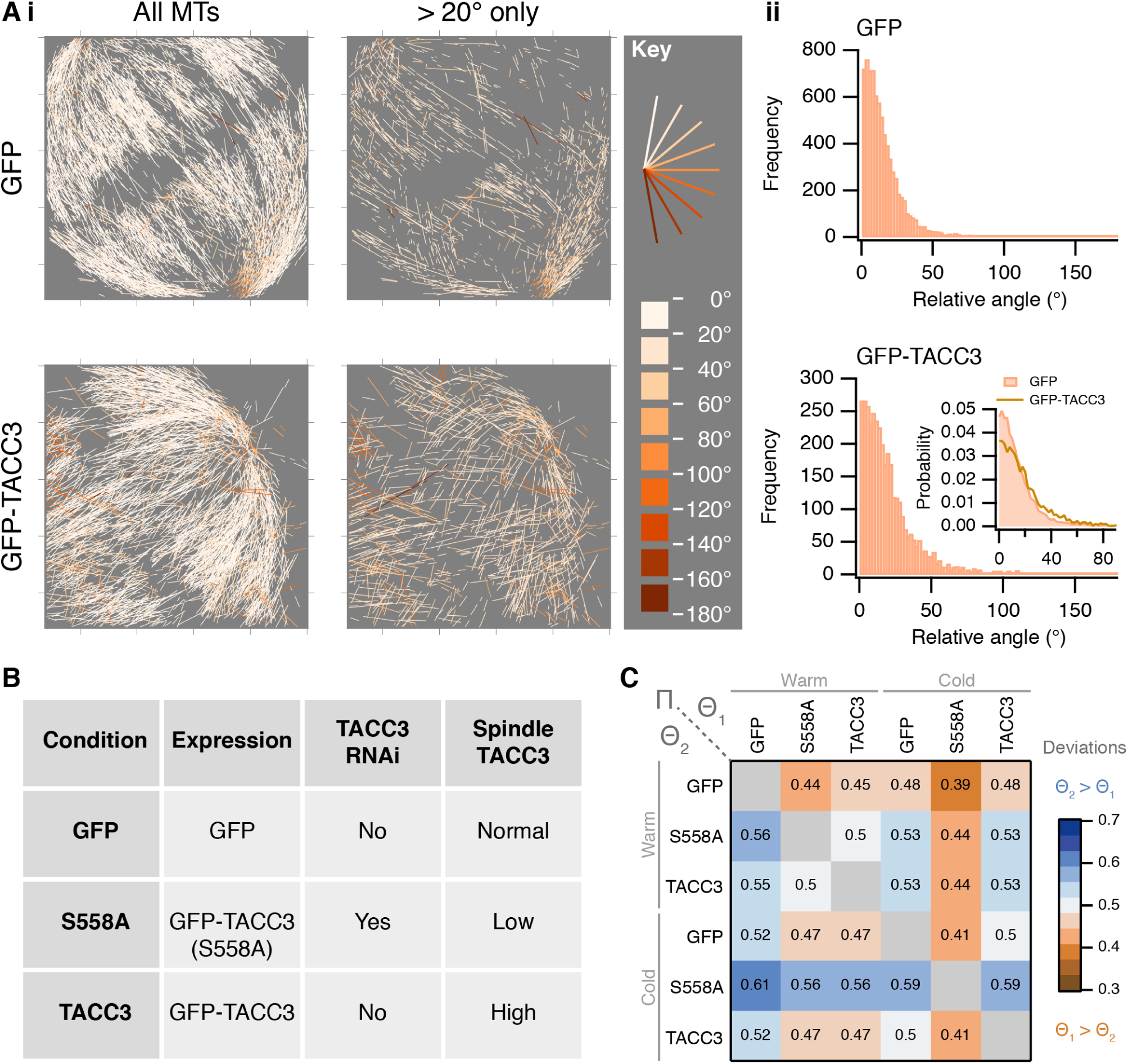
A statistic for microtubule order in mitotic spindles. (**A**) Two examples of angle analysis. (**i**) MTs from a control cell and a cell overexpressing GFP-TACC3 are shown colored according to their deviation from the respective model spindle. In the right panel, MTs with deviations less than 20.0° have been removed for clarity. (**ii**) Histogram of angle deviations for the spindles. (**B**) Summary of cellular conditions analyzed under warm and cold conditions. (**C**) Heat map to show Π(Θ_1_, Θ_2_) for several conditions as indicated. Π is a comparison of angles from two data sets: values greater than 0.5 indicate that Θ_2_ MTs deviate more from the ideal data than those in Θ_1_ (see Methods section 4.4.5 for further details). *N*_segments_ = 5031 − 23240, from *N*_cell_ = 1 − 4.

**Figure 5:**
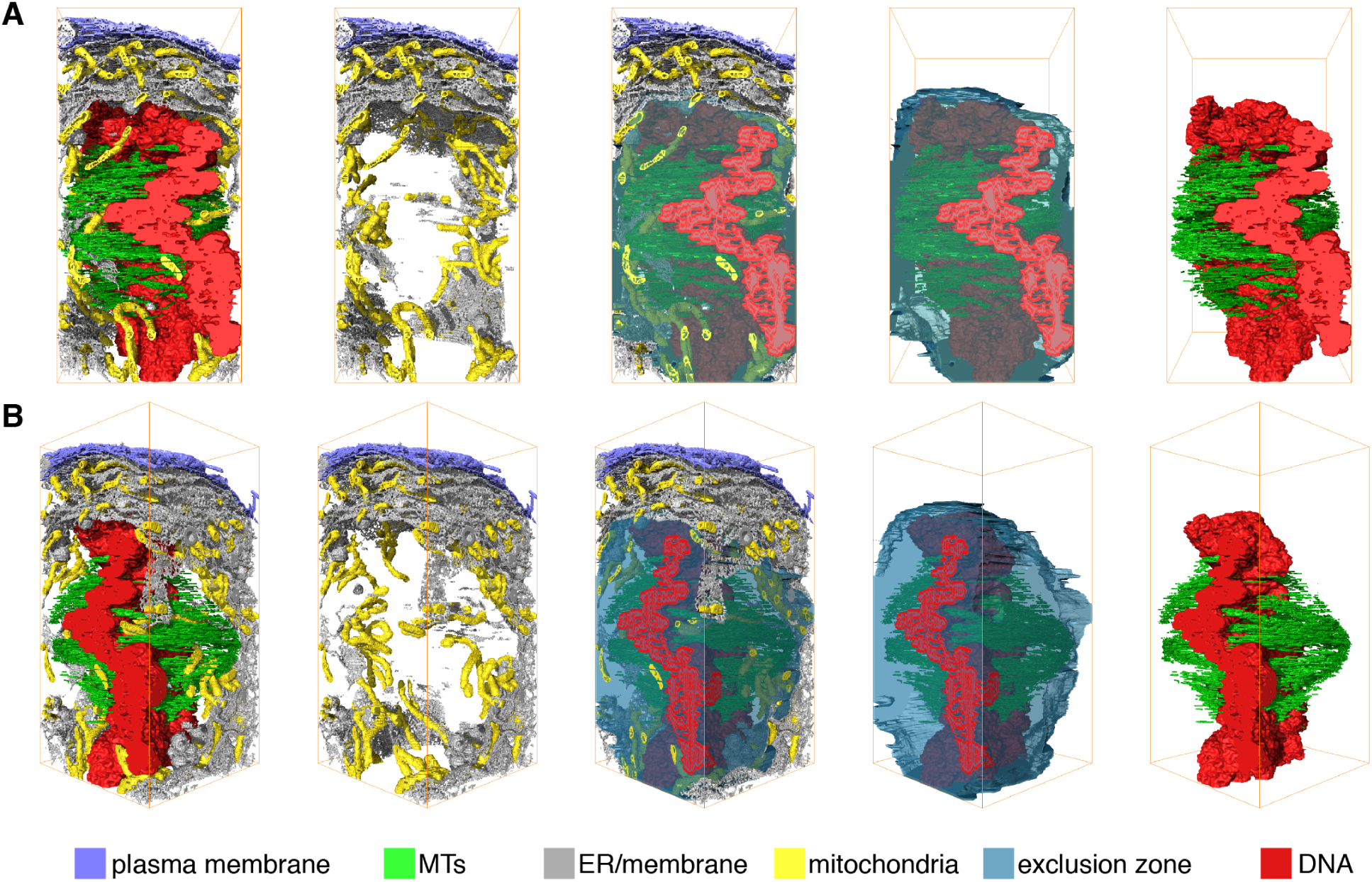
The spindle exists in an “exclusion zone”. 3D rendered model of a mitotic cell at metaphase to show the organization of membranes relative to the mitotic spindle. Subcellular structures that are rendered in 3D: chromosomes (red), mitotic spindle microtubules (green), mitochondria (yellow), plasma membrane (purple), and endoplasmic reticulum and other membranes (gray). An “exclusion zone” is modeled as a translucent blue bubble. The view in **A** is rotated 45° about the Z-axis to give the view in **B**. **A** larger version of this figure with more rotations is shown in Supplementary Figure 1.

**Figure 6:**
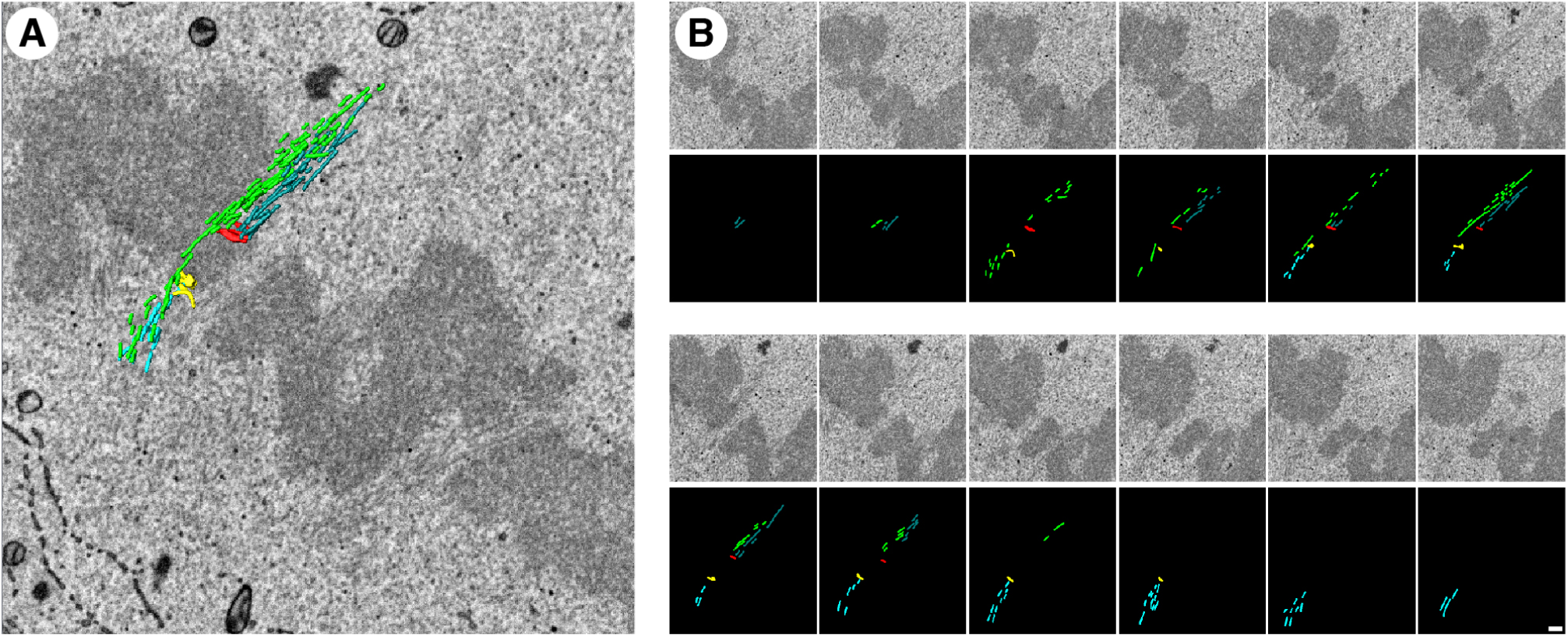
Example of a putative bridging fiber. (**A**) A single SBF-SEM image is shown of the area of interest, with a 3D model of microtubules and kinetochores overlaid. Opposing sister kinetochores (red and yellow) are connected to K-fibers (dark blue and light blue, respectively). (**B**) Sequential SEM images (above) with segmentation (below) that make up the image shown in (**A**). Each image is separated by 60nm in the z direction. Note that, in this example the bridging fiber runs on the exterior of the spindle not the interior as predicted (Kajtez et al., 2016). A movie showing putative bridging fibers is shown in Supplementary Movie 3.

## 3 Discussion

SBF-SEM is commonly used to examine cellular and multicellular structures on the scale of tens to hundreds of micrometers. Whether or not this mode of imaging could discern small subcellular structures such as MTs (25nm diam.) in complex networks such as the mammalian mitotic spindle was untested. Our results indicate that it is possible to visualize a large sample of MTs in the mitotic spindle of a human cell. The resulting datasets were complex and required robust analytical tools which were developed and presented as part of this paper. We applied this workflow to test the influence of TACC3 levels at the mitotic spindle on the organization of MTs on a whole-spindle scale.

The influence of TACC3 on mitotic spindle structure has previously been examined using two extreme methods. In the first, individual high magnification views of subsections of the spindle have been viewed by electron tomography (Nixon et al., 2015). Second, low spatial resolution views of whole spindles were captured with light microscopy (Hood et al., 2013; Gergely et al., 2000). What was missing was a technique which combines the advantages of both methods to visualize MT order on the scale of the whole mitotic spindle. The present study shows that SBF-SEM can fill this resolution gap – at an appropriate scale – to answer questions about MT organization. Moreover, the automation of the image capture and data processing mean that it is practical to analyze spindles in this way and that it may be employed by groups without advanced electron microscopy knowledge.

The analytical workflow allowed us to assess MT density and MT order. While it is possible to approximate the volume of MTs in cells using light microscopy, the higher resolution of SBF-SEM provides more precise quantification of the MT volume in a mitotic cell. Determining the order of MTs in a dense array is not currently possible by light microscopy and the advantage of SBF-SEM here is therefore clear. The simple description that we developed was based on comparison with an idealized spindle. We found that ellipsoids were a good model for angular comparison, out-performing other geometric shapes and comparison regimes based on near-neighbors or on the spindle axis.

Our data indicate that too much TACC3 or too little a affects MT order in mitotic spindles. Previous work has shown that mitotic progression is slowed in cells overexpressing TACC3 or in cells depleted of TACC3 using RNAi (Nixon et al., 2015; Gergely et al., 2003; Lin et al., 2010). The current picture is that the major function of TACC3 in mitotic spindles at metaphase is as part of a TACC3–ch-TOG–clathrin complex that is assembled by Aurora-A phosphorylation, rather than as a MT +tip protein. The TACC3–ch-TOG–clathrin complex is an important component of the “mesh” which is thought to maintain the integrity of the K-fiber MTs. The present data indicate that the sensitivity of mitosis to TACC3 levels is because of changes in the composition of the mesh which alter K-fiber structure and organization; it strongly suggests that these changes prevent the chromosome movements required for high-fidelity mitosis. However, we cannot exclude the possibility that other functions of TACC3 contribute to MT order or to mitotic progression.

These 3D views of mitotic cells also allowed us to visualize the spindle exclusion zone and putative “bridging fibers” of the mitotic spindle. Clearly distinguishing *bona fide* bridging fibers from other microtubules is challenging, even at this resolution. Some examples presented here clearly look like bridging fibers which run in parallel to the respective sister K-fibers (Kajtez et al., 2016; Tolic and Pavin, 2016). Others highlighted in Supplementary Video 3 were less clear. If these structures are not bridging fibers, they could be interpolar MTs, or K-fiber MTs which are laterally or merotelically attached to kinetochores (Cimini et al., 2001). A formal test to determine if these MT bundles are associated with the sister K-fibers will require TEM analysis.

Analysis of MT order by SBF-SEM requires operating at the resolution limit of the instrument. We optimized the staining of the sample, section depth, the magnification and accelerating voltage/beam energy to be able to visualize MTs. This protocol allowed us to sample a large fraction of MTs in the spindle, but fell short of a complete view of the mitotic spindle for three reasons. First, the spindle volume was incomplete since the magnification required to see MTs resulted in an XY area of 9.2 x 9.2 µm. Second, while MTs can be observed, the low number of MTs per K-fiber and the lack of astral MTs (which exist as single MTs) indicate that we were unable to consistently achieve single MT resolution. Third, the section depth of 60nm resulted in missing MT information. The step size matched the beam penetration depth, because heavily stained objects such as membranes could be accurately followed with no under- or over-sampling. However, thin objects such as MTs that are weakly stained appear discontinuous presumably because they are only visible at the upper surface of the block. The spindle exclusion zone assists MT visualization because the organelle-free zone does not interfere with MT detection and segmentation. For these reasons, one caveat associated with the method presented here is that the analysis is performed on only a subset of the MTs in the mitotic spindle. We believe that this subset is a good representation of the mitotic spindle. It was sufficient to show differences in MT density and changes in MT order in the mitotic spindle. Improvements in detection may allow us to improve our coverage, but the protocol presented here is sufficient for large-scale analysis of MT order in human cells.

## 4 Materials and Methods

### 4.1 Cell biology

HeLa cells which overexpress TACC3 upon doxycycline induction were described previously (Nixon et al., 2015). Cells were maintained in Dulbecco’s Modified Eagle’s Medium (DMEM) plus 10% fetal bovine serum (FBS) and 1% penicillin/streptomycin, in a humidified incubator at 37°C and 5% CO2. The cell culture medium was further supplemented with G418 (300 µg ml^−1^) for the parental HeLa TetOn cells, and with G418 and Hygromycin B (200 µg ml^−1^) for the HeLa TetOn cells stably transfected with the TACC3 plasmid. Glioblastoma cell lines were cultured using Neurobasal medium, supplemented with B27, N2, human fibroblast growth factor (40 ng ml ^−1^) and human epidermal growth factor (40 ng ml^−1^).

For experiments to express GFP-TACC3(S558A), we used a plasmid (pBrain-GFP-TACC3(S558A)KDP-shTACC3) which knocks down the expression of endogenous TACC3 and simultaneously expresses GFP-TACC3(S558A) which is refractory to the shRNA (Gutiérrez-Caballero et al., 2015). Cells were transfected using GeneJuice, according to manufacturer's instructions.

Three HeLa conditions were studied: first, parental HeLa TetOn cells transiently transfected with pEGFP-C1 only as a control; second, the inducible GFP-TACC3 HeLa TetOn stable cell line; third, parental HeLa TetOn cells transiently transfected with pBrain-GFP-TACC3(S558A)KDP-shTACC3 construct. Cells were synchronized by thymidine-Ro3306 treatment (Nixon et al., 2015) and grown on gridded dishes (MatTek) to allow correlation between light and electron microscopy. For cold treatment of mitotic cells, cells were imaged to ensure the cells were at metaphase, the growth media removed and replaced with ice-cold growth media and incubated on ice for 10 min. Media was then replaced with cold fixative.

### 4.2 Electron Microscopy

Fluorescence imaging was done using a Nikon Eclipse Ti-U microscope with standard filter sets for visualization of GFP. Light and epifluorescence micrographs were acquired using a Photometrics Myo camera, 20x air and 60x oil objectives, and NIS Elements acquisition software. Cells were imaged in a 37°C temperature-controlled chamber (OKOlab). Chemical fixation, processing and sectioning for TEM, was done as described previously (Booth et al., 2013). To prepare samples for SBF-SEM, the cells were also processed in a correlative manner to ensure only metaphase cells expressing the protein of interest were chosen. Processing was as described with the following extra processing steps to better visualize ultrastructural components of the cell by SEM. Formaldehyde (0.5%) and glutaraldehyde (3%) fixative solution was prepared in phosphate buffer (0.1 mol l^−1^), with 0.1% tannic acid and 3% sucrose. Two osmication steps: first, a 1 h reduced osmium step where 2% OsO_4_ was prepared in 1.5% potassium ferrocyanide solution, in phosphate buffer. Five 3 min ddH_2_O washes, before 0.1% thiocarbohydrazide (TCH) was added for 30 min at 60 °C as a mordant. Five 3 min ddH_2_O washes. Second, OsO_4_ step (2% in ddH_2_O) was applied for 30 min. The cells were again washed in water as before, then stained overnight using 1% uranyl acetate in ddH_2_O at 4 °C. The following day, water washes were repeated before incubating with lead aspartate at 60 °C for 30 min. The cells were then dehydrated using graded ethanol. Cells were infiltrated using Epon 812 Hard premixed resin kit (TAAB). Once fully polymerized (48 h at 60 °C), dish and coverslip were removed and the cell located. This area was then excised using a junior hacksaw and razor blades to leave an approximately 2 mm^3^ piece of resin containing the cell. This cube of resin was super-glued onto special steel pins which fit into the chuck of the 3View platform (Agar Scientific). The area around the cell was fine-trimmed using fresh glass knives, leaving as little resin as possible around the cell of interest to reduce interference produced by charging of the resin surface during SEM imaging. This produced an approximately 100 µm block face. Due to the design of the 3View stage and ultramicrotome, the majority of the 2 mm^3^ block needed to be trimmed away to avoid excess resin catching on the knife blade or holder. This produced an extremely delicate block face so special care was needed to prevent damage during manipulation. Once trimmed, the block was coated with a silver paint using an eyelash, before a 15nm gold/palladium coating was evaporated onto the block surface, again to reduce surface charging (Quorum Technologies, East Sussex, UK). Once dry, the pin/block was mounted into the stage and aligned manually with the knife-edge. When aligned, the 3View was allowed to cut-and-image the block surface, with 100nm z-slices, initially optimizing the contrast and magni cation for that particular block, before leaving the platform to run until the cell was fully sectioned and imaged. A 768 x 768 pixel image, 60nm sections, 60 µs pixel dwell time, 2:1 keV, and 21000x magnification (12nm pixel size, 9.216 x 9:216 µm) was optimal for imaging the spindle zone. These conditions needed to be fine-tuned for the individual sample, and were a compromise between reduced levels of sample charging, magnification and resolution.

### 4.3 Image analysis

All 3D segmentation and surface rendering was performed using `Amira` software (Visualization Sciences Group, FEI). The voxel size in nm was entered upon loading of the le into `Amira`. MTs in SBF-SEM volume stacks were segmented manually in `Amira` by tracing all the MTs visible in each slice throughout the entire volume, this segmentation was rendered to generate a 3D surface. In addition, where membranes, chromosomes, kinetochores and centrioles were visible, these were segmented in a semi-automated manner, using pixel density to segment objects in 3D. The segmentation les (*.`am` les) were processed in `ImageJ/FIJI` to create binary images which could be fed into `Igor Pro 7.0.` Again code for analysis was custom-written in `Igor Pro 7.0` and can be found at https://github.com/quantixed/VolumeFinder. For volume analysis, the image stacks are interpreted and pixels above threshold are counted for each slice of the stack. A 3D convex hull is drawn around all points to measure the volume for normalization. For ellipsoid comparison, described in detail below, the images are skeletonized in `ImageJ/FIJI`. This process converts each MT segment into a 1 pixel thick object. Each object is read by `Igor` and converted to a unique 2D vector by fitting a line *y* = *ax* + *b* to the *xy* coordinates of the object. All objects lie in the plane of the section/image and so a 2D vector is sufficient. The resulting straight lines are used for mathematical modeling. The modeling and comparison was done in `Igor Pro 7.0` for code, see https://github.com/quantixed/VolumeFinder. To ensure reproducibility and for error-checking the modeling and analysis was independently rewritten in `R` using the *xyz* coordinates of straight lines as an input.

### 4.4 Mathematical modeling

A more detailed description follows; in summary, straight lines (representing MT segments from the SBF-SEM dataset) were compared to computer-generated counterparts that were placed in the ideal location. All straight lines were translated and rotated such that the spindle was vertical with the poles aligned on the z-axis and the center of the spindle at the origin. The midpoint of each straight line was used to make an ellipsoid that passes through this point and through both spindle poles. The tangent of a section through this ellipsoid at the midpoint was used for comparison with the original straight line. This was done after putting the straight lines and the idealized counterparts back in their original location, and projecting both segments onto a single xy-plane. We developed this analysis method because the spindle is a fusiform structure, and therefore comparison of each MT segment to the spindle axis gave a broad range of deviations that made comparisons between conditions impossible. Similarly, angle comparisons between each MT segment and its neighboring segments was also highly variable. The calculations rely on simple geometric principles and make use of specified locations of the two spindle poles. This is possible because the force balance in the metaphase spindle ensures a symmetrical arrangement: the chromosomes are situated approximately at the equator, and the spindle poles are equidistant from the metaphase plate.

#### 4.4.1 Observed lines

We observe a collection of straight lines, *i* ∈ {1, 2, …, *n*}, denoted 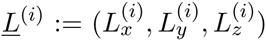. Each line may be summarized by its two endpoints; *<u>L</u>*(0) and *<u>L</u>*^(*i*)^(1), switching the labels of endpoints *<u>L</u>*^(*i*)^(0) and *<u>L</u>*^(*i*)^(1) results in the same line object. Section 4.4.3 justifies a particular choice of *<u>L</u>*^(*i*)^(0) and *<u>L</u>*^(*i*)^(1).

Line locations and orientations are determined from a stack of images collected at increasing depth through the cell with each line segment being extracted from a single image within this stack as described in Section 4.3. The co-ordinate axes are aligned such that the image planes are perpendicular to the *z*-direction.

#### 4.4.2 Elliptical model

We wish to compare observed lines to paths lying on the surface of an ellipsoid model. In fact, we restrict ourselves to spheroids (i.e. ellipsoids with rotational symmetry about the *z* axis) because we do not expect there to be any promotion/restriction to MT growth in either the *x*- or *y*-directions. The surface of such a spheroid centered at the origin contains all vectors of the form 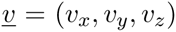 such that

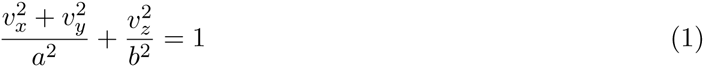

Proposed paths are geodesic curves on the surface of an ellipsoid described by (1) which pass through the points (0, 0, *b*) and (0, 0, −*b*). Fixing the value of *b* ensures that each path passes through a common pair of points, which model the mitotic spindle poles. This modeling is justified biologically since the force-balance of the metaphase spindle ensures that the poles are approximately equidistant from the chromosomes at the equator. The remaining principal semi-axis length, *a*, is allowed to vary between paths, with *a*^(*^i^*)^ used to denote the value associated with the line indexed by *i*, ensuring that every point in the plane *z* = 0 is associated with the unique path that passes through it, representing the unique proposed trajectory of the MT which attaches to a theoretical chromosome at that location.

The formula given in (1) describes a spheroid centered at the origin and aligned with the co-ordinate axes. However, while we expect our observed lines to be related these trajectories, we do not expect them to be centered and aligned. It is possible to transform the model to match the location and alignment of the observed data. We choose instead to transform the observed data for later ease of interpretation and presentation.

#### 4.4.3 Translation and rotation

Each observed line-segment is first translated so that the origin of the spheroid whose tangent-plane contains it is centered at the origin and then rotated so that this spheroid is aligned with the coordinate axes. This process begins by identification of two fixed points, *<u>p</u>*^(1)^ and *<u>p</u>*^(2)^, which after translation and rotation will become (0, 0, *b*) and (0, 0, −*b*) respectively. The location of these fixed points can be determined directly from the images themselves, in the case where the centrosomes are visible. Where centrosomes are not directly visible, they were inferred using the orientation of observed lines and the assumption that *<u>p</u>*^(1)^ and *<u>p</u>*^(2)^ are the spindle poles from which all MTs should originate.

The translation shifts the center of the straight line joining *<u>p</u>*^(1)^ and *<u>p</u>*^(2)^ to the origin by subtracting the centre *<u>c</u>* = (*<u>p</u>*^(1)^ + *<u>p</u>*^(2)^)/2 of the spheroid from each observation. Rotation about the line through the origin which is perpendicular to both the *z*-axis and the line-segment connecting *p*^(1)^ to *<u>p</u>*^(2)^ by the angle required to align the image of that line-segment with the *z*-axis is then carried out. We allow *<u>λ</u>*^(*i*)^ to denote the shifted and rotated line segment obtained by transforming *<u>L</u>*^(*i*)^ in this way, with end points *λ*^(*i*)^(0) and *<u>λ</u>*^(*i*)^(1).

Under our model, we expect observed MT trajectories to initiate at (0, 0, ±*b*) and to terminate at the plane *z* = 0. We orient each trajectory so that it is consistent with this. This allows us to specify that *<u>λ</u>*^(*i*)^(0) is the end of each line segment closest to a fixed point and *λ*^(*i*)^(1) the end closest to the plane which models the metaphase plate.

Our interest is focused on those MTs leading from the spindle pole to the metaphase plate and so our analysis is restricted to observed lines for which the *z*-coordinate lies always in [−*b*, +*b*].

#### 4.4.4 Model directions

We wish to compare the direction of observed lines <u>*λ*</u>^(*i*)^, conveniently denoted <u>*δ*</u>^(*i*)^ = *<u>λ</u>*^(*i*)^(1) – <u>*λ*</u>^(*i*)^(0), with those consistent with the spheroid model. The first step is to determine the tangent to the geodesic path which passes through the fixed points (0, 0, *b*), (0, 0, −*b*) and the midpoint of the observed line *<u>λ</u>*^(*i*)^, given by *<u>μ</u>* ^(*i*)^=(*<u>λ</u>*^(*i*)^(0) + *<u>λ</u>*^(*i*)^(1))/2, on the surface of the spheroid.

The unique spheroid which passes through *<u>μ</u>*^(*i*)^ and both (0, 0, *b*) and (0, 0, − *b*) is determined by the value of *a*^(*i*)^ given by

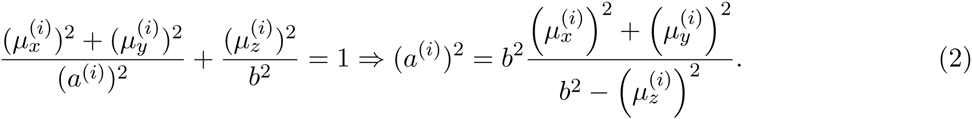

The unique tangent to the path at that point is ascertained by standard geometric arguments and is given by:

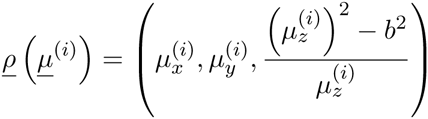

Comparison could be made between <u>*ρ*</u>(*<u>μ</u>*^(*i*)^) and *<u>δ</u>*^(*i*)^. However, the earlier remark that all observed lines *<u>L</u>*^(*i*)^ lie within parallel planes means that even if the MT trajectories exactly follow the ellipsoid model proposed, *<u>δ</u>*^(*i*)^ and <u>*ρ*</u>(*<u>μ</u>*(*i*)) will not be identical. This discrepancy may be removed by comparing *<u>δ</u>*^(*i*)^ to 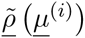, the projection of *<u>ρ</u>*(*<u>μ</u>*^<u>^(^</u>*<u>^i^</u>*<u>^)^</u>^) into the observation planes.

In the original orientation, lines *<u>L</u>*^(*i*)^(*t*) lie in one of the image planes which are perpendicular to the *z*-axis and therefore perpendicular to the vector 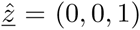. After translation and rotation the lines *<u>λ</u>*^(*i*)^ no longer lie in planes perpendicular to 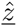, rather they lie in planes perpendicular to 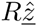 where *R* is the matrix which characterizes the rotation component of the transformation applied to the original line (in the canonical Euclidean basis). As a result, the projection of proposed directions *<u>ρ</u>*(*<u>μ</u>*^(*i*)^) into the planes, 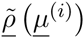, is given by

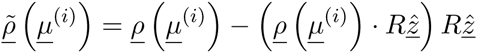

#### 4.4.5 Model discrepancy and comparison

Angles, denoted by *θ*^(*i*)^, between observed directions, *<u>δ</u>*^(*i*)^, and model directions, 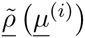, can be calculated as a measure of the discrepancy between observations and the proposed model:

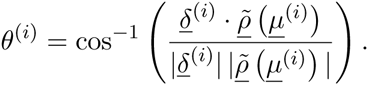

The set of deviations Θ= {*θ*^(1)^, *θ*^(2)^; …, *θ*^(*n*)^ can be visualized using a histogram as shown in Figure 4. Note, that systematic variation in the reported angle deviations based on location in the spindle or due to segment length was looked for and no evidence was found.

Analyzing two sets of observations produces two sets of angles, Θ_1_ and Θ_2_, which may be compared to suggest whether one observation deviates more from the proposed ellipsoid model than the other. We allow Π(Θ_1_, Θ_2_) to denote the proportion of all pairs (*θ*, *θ*) ∈ Θ _1 ×_Θ_2_ in which *θ* < *υ*:

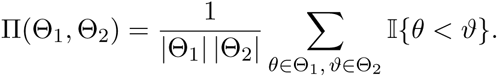

where 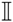 denotes the indicator function which takes the value 1 if its argument is true and 0 otherwise. The summary Π(Θ_1_, Θ_2_) takes values in [0, 1]: Values of (Θ_1_, Θ_2_) close to 0.5 suggest that the model fits each collection of observations equally well, values below 0.5 suggest that deviations from the model are generally greater for the first set of observations while values greater than 0.5 suggest that deviations from the model are generally greater for the second set of observations. Comparison using Π is preferred to comparison of means as it has greater robustness.

## 5 Manuscript information

### 5.1 Acknowledgments

We thank Heiko Wurdak for the gift of glioblastoma cell lines. This work was supported by a North West Cancer Project grant (CR928) and a Senior Cancer Research UK Fellowship (C25425/A15182) to SJR.

### 5.2 Author contributions

FMN: did the experimental work, segmentation, rendering and analysis, TRH: mathematical modeling, wrote R code, GS: additional segmentation and rendering, AJB: assisted with SBF-SEM, JAB: mathematical modeling and co-supervision of TRH, AMJ: mathematical modeling and co-supervision of TRH, IAP: co-supervised FMN, SJR: analyzed data, wrote Igor code, wrote the paper, co-supervision of FMN.

The authors declare no conflict of interest

## Supplementary Files

**Supplementary Movie 1.**
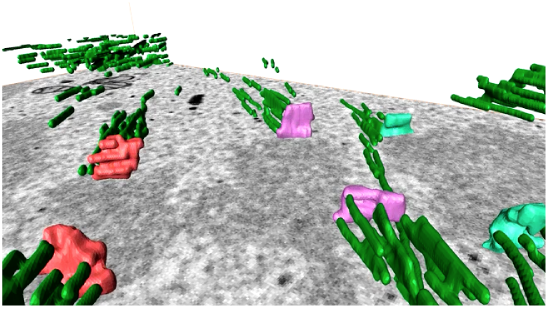
Movie to show 3D rendering of a SBF-SEM dataset from a HeLa cell at metaphase. MTs (green) are shown together with chromosomes (red) and kinetochores, where visible are shown are colored patches. Same colors indicate the sister pairs.

**Supplementary Movie 2.**
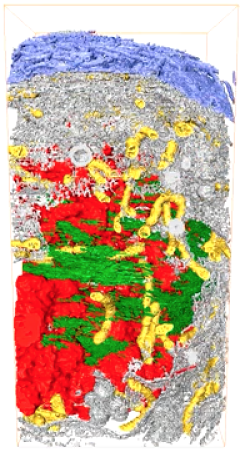
Spindle exclusion zone. A movie of 3D rendering of an SBF-SEM dataset from a HeLa cell at metaphase. MTs (green) are shown together with chromosomes (red) and membranes are colored as described in Figure 5. The exclusion zone is modeled as a translucent blue bubble.

**Supplementary Movie 3.**
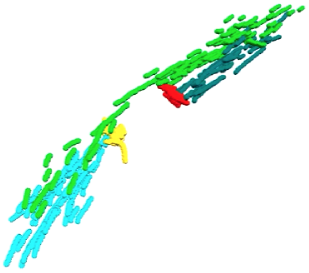
Bridging fibers. A movie of 3D rendering of an SBF-SEM dataset from a HeLa cell at metaphase to accompany Figure 6.

**Supplementary Figure 1.**
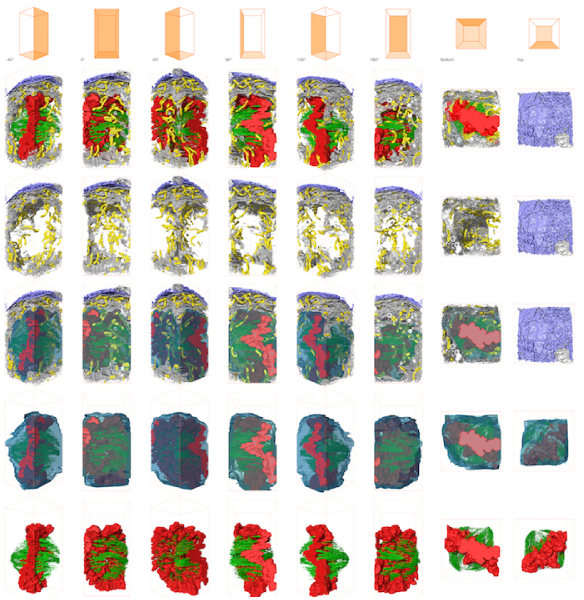
The mitotic spindle exists in a spindle “exclusion zone”. Various views of a 3D rendering of an SBF-SEM dataset from a HeLa cell at metaphase to accompany Figure 5.

